# The connection between the gut mycobiome community of natural black soldier fly populations and their environment

**DOI:** 10.1101/2022.03.28.486161

**Authors:** Tzach Vitenberg, Martin Goldway, Itai Opatovsky

## Abstract

In contrast to interactions between prokaryotic organisms (e.g. bacteria) and insects, an understanding of the interactions between eukaryotic microorganisms (e.g. fungi) and insects is lacking. In this study, black soldier fly larvae were collected from natural populations in household composts. The mycobiome composition of the larval gut and environment was identified using next generation sequencing. The nutrient composition of the larvae’s environment was analyzed. The mycobiome composition was divided into three groups: the first included larvae from Kfar HaHoresh that comprised mostly *Meyerozyma* and were influenced by the fiber and carbohydrate composition of the compost; the second included the compost samples from Kfar HaHoresh and Misgav Am that mainly comprised the Dipodascaceae family and were influenced by the fiber composition of the compost. The third group contained all the remaining compost and larvae samples and mainly comprised the genus *Candida*. *Candida* spp. are the most common in the environment and may thus affect their abundance in the insect gut. However, this species may be adapted to the insect’s gut, providing the insect with a benefit that allows the insect to disperse *Candida* to new resource patches and thereby help increase its abundance in the environment. In cases where dominant species are not present, it seems that the important factor determining fungal composition in the insect gut is its diet. Understanding the metabolic effect of fungi, e.g. *Candida*, on the BSF larvae is still a black box which requires further study.

**IMPORTANCE:** The research analyzed the mycobiome composition of BSF larvae from natural environments. It was found that the most dominant fungi in these communities is the Candida genus, which was found also dominant in the BSF environment (household compost) that differ in its nutrient composition. Understanding the metabolic effect of these fungi on the BSF larvae have the potential to dramatically affect the physiological condition of the insect. Revealing these metabolic interactions may help to increase the rearing efficiency of insects, such as the BSF, which is currently being reared at large scales as an alternative protein source.

## INTRODUCTION

Interactions between microorganisms and insects are known mainly from prokaryotic organisms (e.g. bacteria), particularly in insects with a homogeneous diet (blood, wood, plant sap), in which the microorganisms provide the components absent from the insect’s diet (1–3). However, an understanding of the interactions between eukaryotic microorganisms (e.g. fungi) and insects is lacking. Fungi, particularly yeast and yeast-like symbionts (hereafter, fungi), have high metabolic complexity and abilities and thus they have the potential to dramatically affect the physiological condition of the insect.

In these mutualistic interactions, the fungi can provide nutrients, such as protein, fatty acids, sterols and vitamins, take part in uric acid and pheromone metabolism (4–6), and protect the host from harmful fungi and parasitoids (7), while the insect host enables dispersal of the fungi to new patches (7). However, it is not clear whether the fungal composition of the insect gut is affected by the fungal composition of the insect’s living environment or whether the fungi adapted to the insect’s digestive system are dispersed by the insects to new habitats, thereby becoming more and more common in the environment, and whether this correlation suggests mutualistic interactions between the fungi and their insect host.

To test these questions we used the black soldier fly (BSF) (*Hermetia illucens*) as the model system for fungal-insect interactions. The BSF is native to the tropical and sub-tropical regions of America and is currently found worldwide (8), mostly in dumpsters and compost bins in urban areas (9). The fly is an omnivorous detritivore that completes its lifecycle in decaying organic matter thriving with microorganisms, some of which may be harmful to the BSF, while others may have mutual interactions with it. In contrast to BSF-bacterium interactions, which have been extensively studied in recent years (10–12), knowledge regarding the interactions of BSF with fungi, particularly metabolic interactions, is lacking. A study that tested the mycobiotic composition of the BSF gut under different diets found that the major yeasts spp. are *Geotrichum candidum*, *Candida tropicalis*, *Pichia fermentans*, *P. kluyveri* and *P. kudriavzevii* (13). However, it is still not known whether the species composition is affected by their density in the environment or whether they are being selected by the BSF. Therefore, we studied natural populations of BSF from Israel, tested the effect of the environmental nutritional conditions on fungal community composition and examined the correlation between fungal composition in the insect’s environment and in its gut.

## RESULTS

### Compost composition

Two sites, KH and M, had the highest carbohydrate content (X^2^_df=5_=12.6, p=0.03) and the lowest fat content (X^2^_df=5_=15.8, p=0.007). Protein content was highest in S and lowest in M (X^2^_df=5_=16.6, p=0.005). Fiber content was highest in KH but not significantly higher than in MA and T (X^2^_df=5_=12.4, p=0.03). Mineral content was highest in M (X^2^_df=5_=16.7, p=0.005) (Table 1).

**Table 1:**
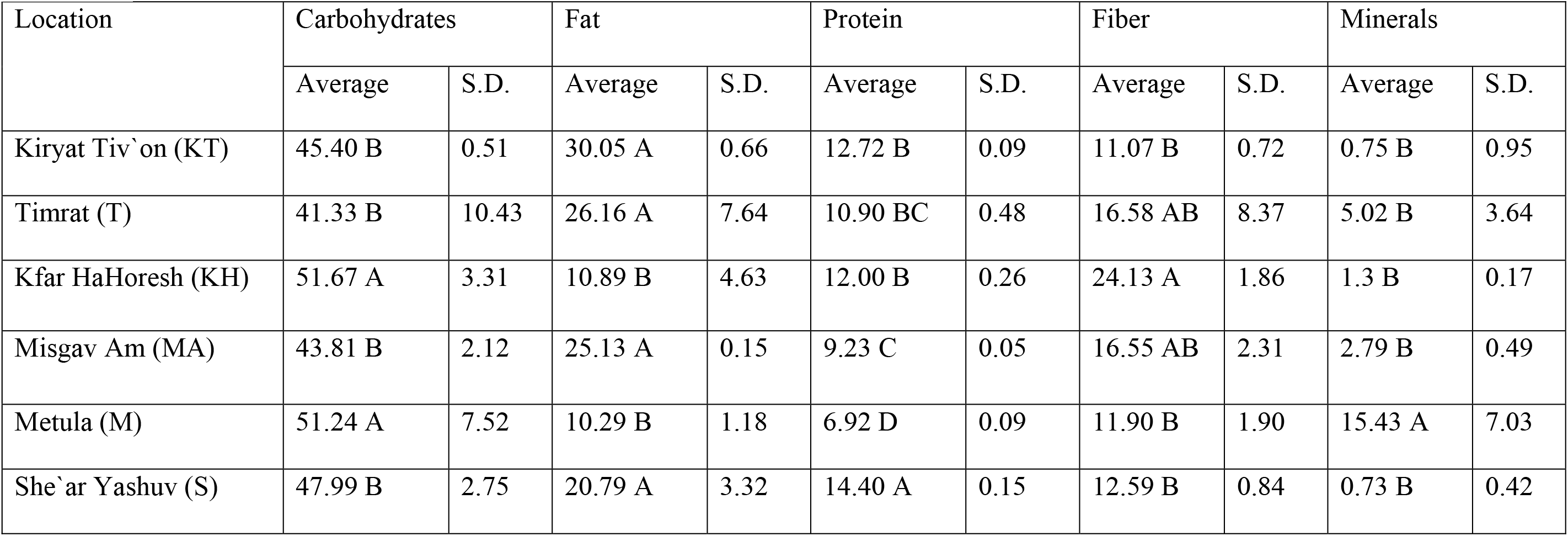
Carbohydrates, fat, protein, fiber and mineral composition of household composts from six locations. Capital letters represent significant differences among the locations (p<0.05).

### Next-Generation Sequencing (NGS) of the mycobiome in the BSF gut and the substrate

In total, 196 genera were identified in the NGS analysis (Supplementary Table 1). The first two PCA axes explained 52% and 23% of the variance, respectively. The analysis divided the locations into three groups: 1) larval samples from KH, dominated by *Meyerozyma*, 2) compost samples from KH and larval and compost samples from MA, dominated by Dipodascaceae, and 3) all remaining compost and larval samples, dominated by *Candida* (Figure 1).

**Figure 1:**
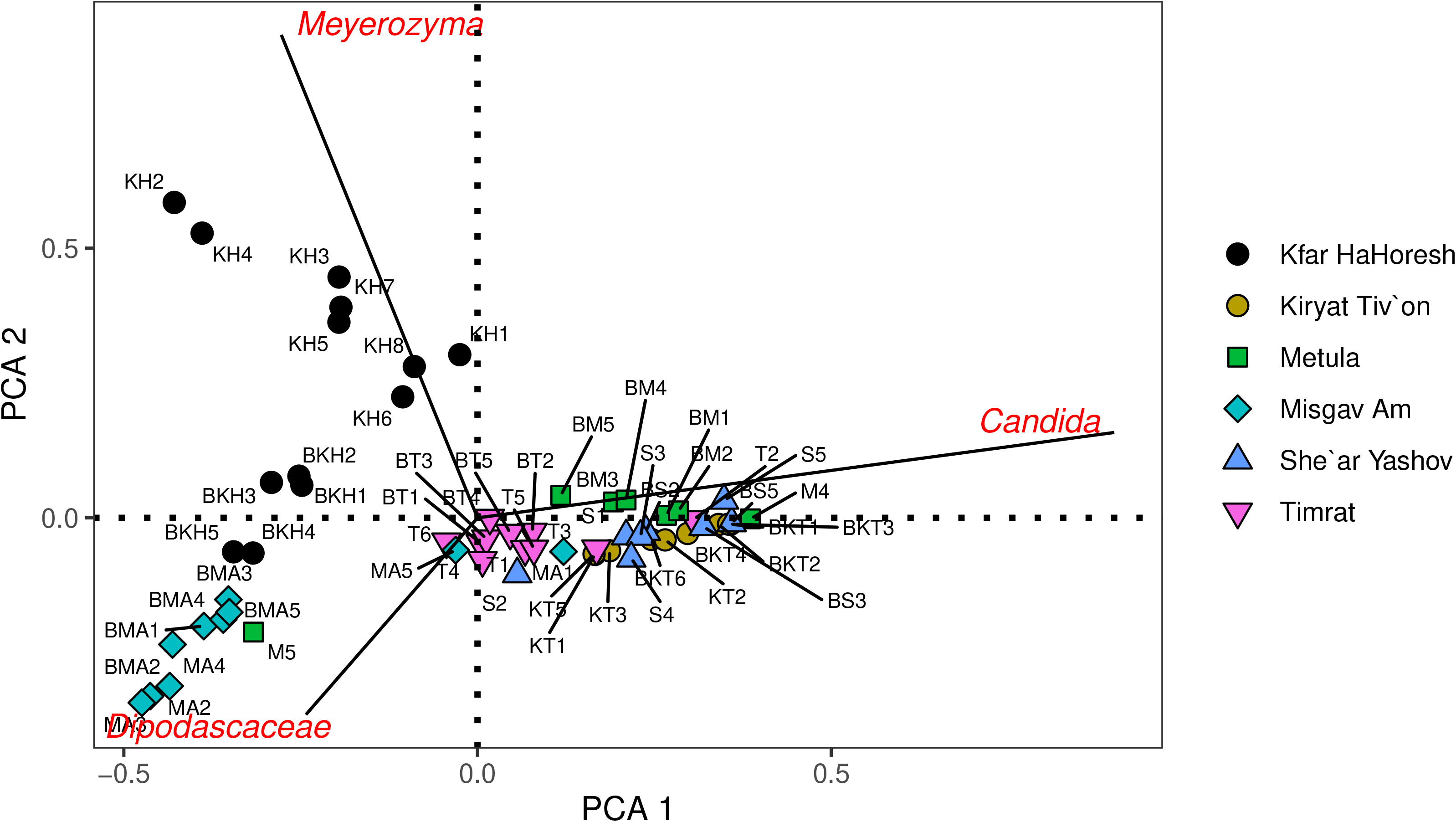
PCA of the fungal composition in the different locations and samples: KH – Kfar HaHoresh larvae; BKH - Kfar HaHoresh compost; KT - Kiryat Tiv’on larvae; BKT - Kiryat Tiv’on compost; T – Timrat larvae; BT – Timrat compost; MA – Misgav Am larvae; BMA - Misgav Am compost; M – Metula larvae; BM – Metula compost; S - She’ar Yashuv larvae; BS - She’ar Yashuv compost. The fungal genera and families that dominated the community composition are marked in red.

To more closely examine the PCA results, we compared the fungal composition of the compost and the larval gut at the different locations (>5% abundance; Figure 2A, B; Supplementary Table 1). The first group included seven fungal genera and families with >5% abundance (*Meyerozyma*, *Candida*, *Gibberella*, Dipodascaceae*, Dipodascus*, *Issatchenkia* and *Arthrographis*) with dominant abundance of *Meyerozyma* and *Candida* (52% and 36%, respectively). The second group, which included the compost and larval samples from MA and compost samples from KH, had 10, 9 and 7 abundant genera and families, respectively, which were more equally distributed without any dominant genus. The MA samples were dominated by Dipodascaceae, with 15% abundance in the compost and 33% abundance in the larvae, while the KH samples were dominated by *Gibberella*, with 27% abundance (Figure 2A; Supplementary Table 1). The third group was dominated by *Candida spp.* (abundance in BKT: 76%; KT: 60%; BT: 46%; T: 55%; BM: 68%; M: 50%; BS: 76% and S: 68%; Figure 2A, B and Supplementary Table 1).

**Figure 2:**
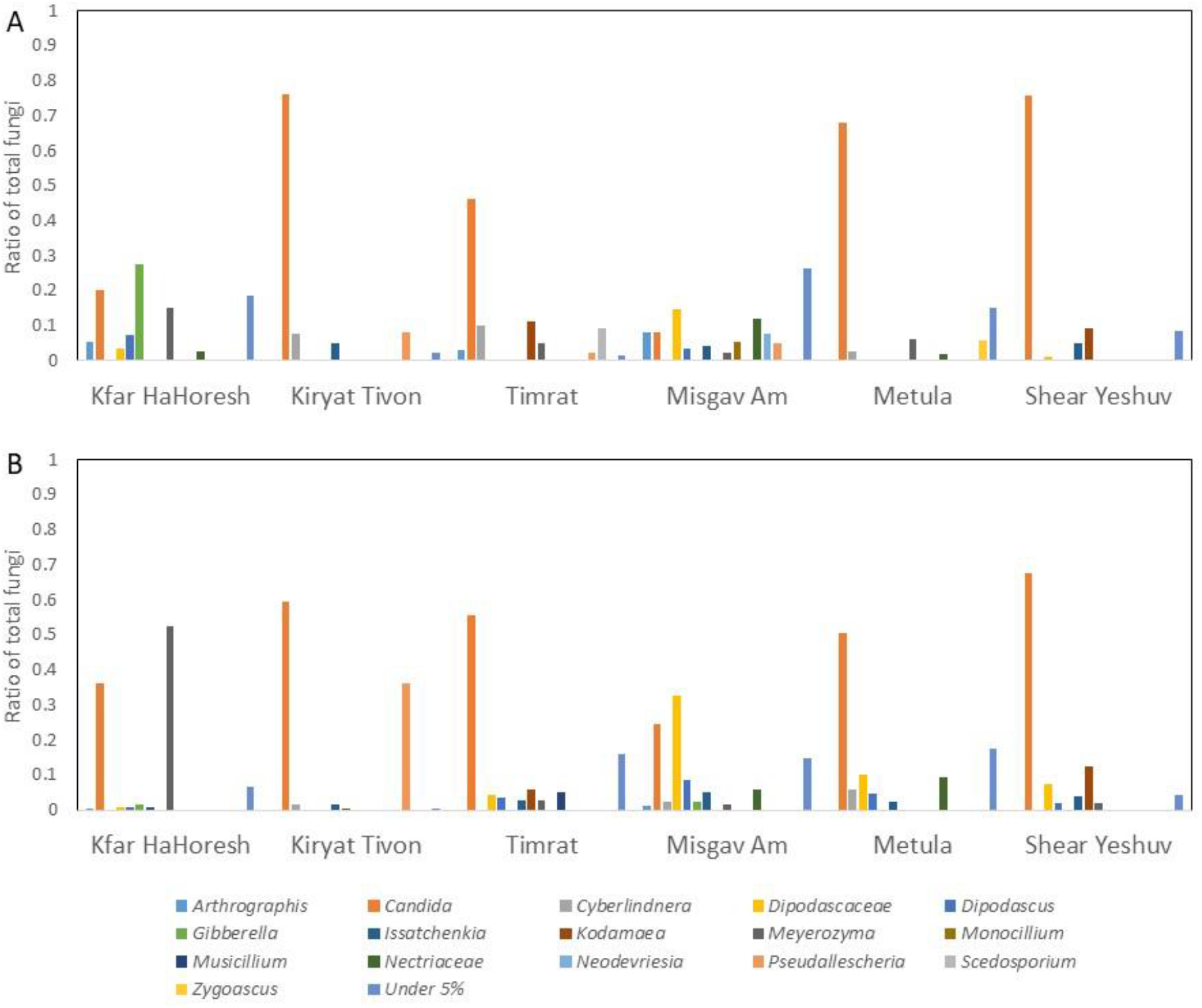
Proportion of each of the main fungal groups (above 5% of abundance) in the compost (A) and insect gut (B) from the different locations.

The first two RDA axes of the compost samples explained 73% and 11% of the variance, respectively. The analysis divided the locations into three groups: 1) the fungal composition of the KH samples, comprising mainly *Gibberella* and *Meyerozyma*, was influenced by the fiber composition of the compost; 2) the fungal composition of the MA samples, comprising mainly Dipodascaceae and Nectriaceae, was influenced by the fat and fiber composition of the compost; and 3) the fungal composition of all remaining samples, comprising mainly *Candida spp*., was influenced by the protein and mineral contents of the compost to varying degrees (T was less influenced by nutrient composition; Figure 3).

**Figure 3:**
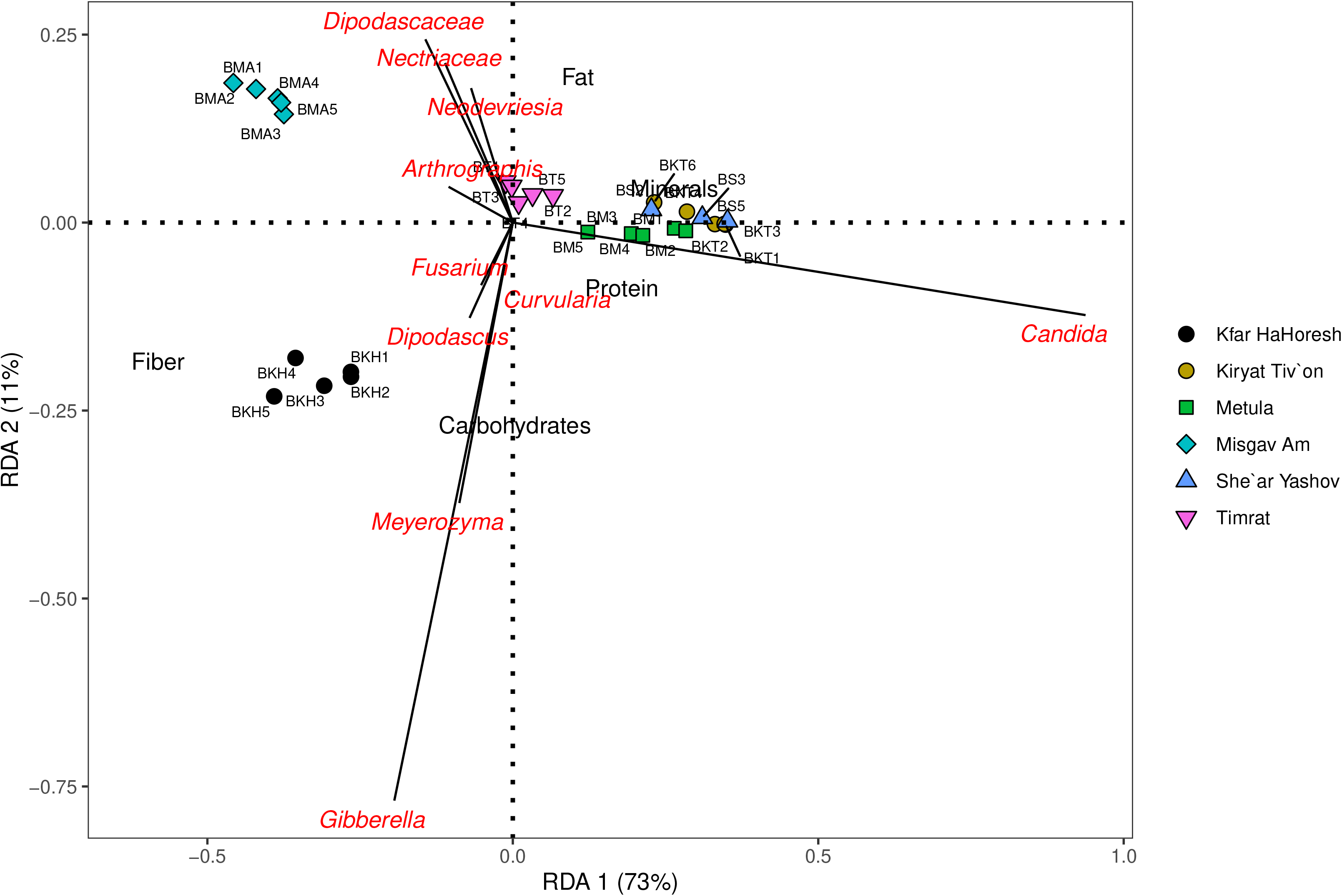
RDA of the fungal composition in the different locations and compost samples: BKH - Kfar HaHoresh; BKT - Kiryat Tiv’on; BT – Timrat; BMA - Misgav Am; BM – Metula; BS - She’ar Yashuv. The fungal genera and families that dominated the community composition are marked in red. The nutrient components of the compost are marked in black.

The first two RDA axes of the larval samples explained 40% and 20% of the variance, respectively. The analysis divided the larval samples into three groups: 1) the fungal composition of the KH samples, comprising mainly *Meyerozyma*, was influenced by the fiber and carbohydrate contents of the compost, 2) the fungal composition of the MA samples, comprising mainly Dipodascaceae, was influenced to a certain extent by the low protein content of the compost, and 3) the remaining larval samples, comprising mainly *Candida spp*., was influenced by the protein and fat contents of the compost to varying degrees (Figure 4).

**Figure 4:**
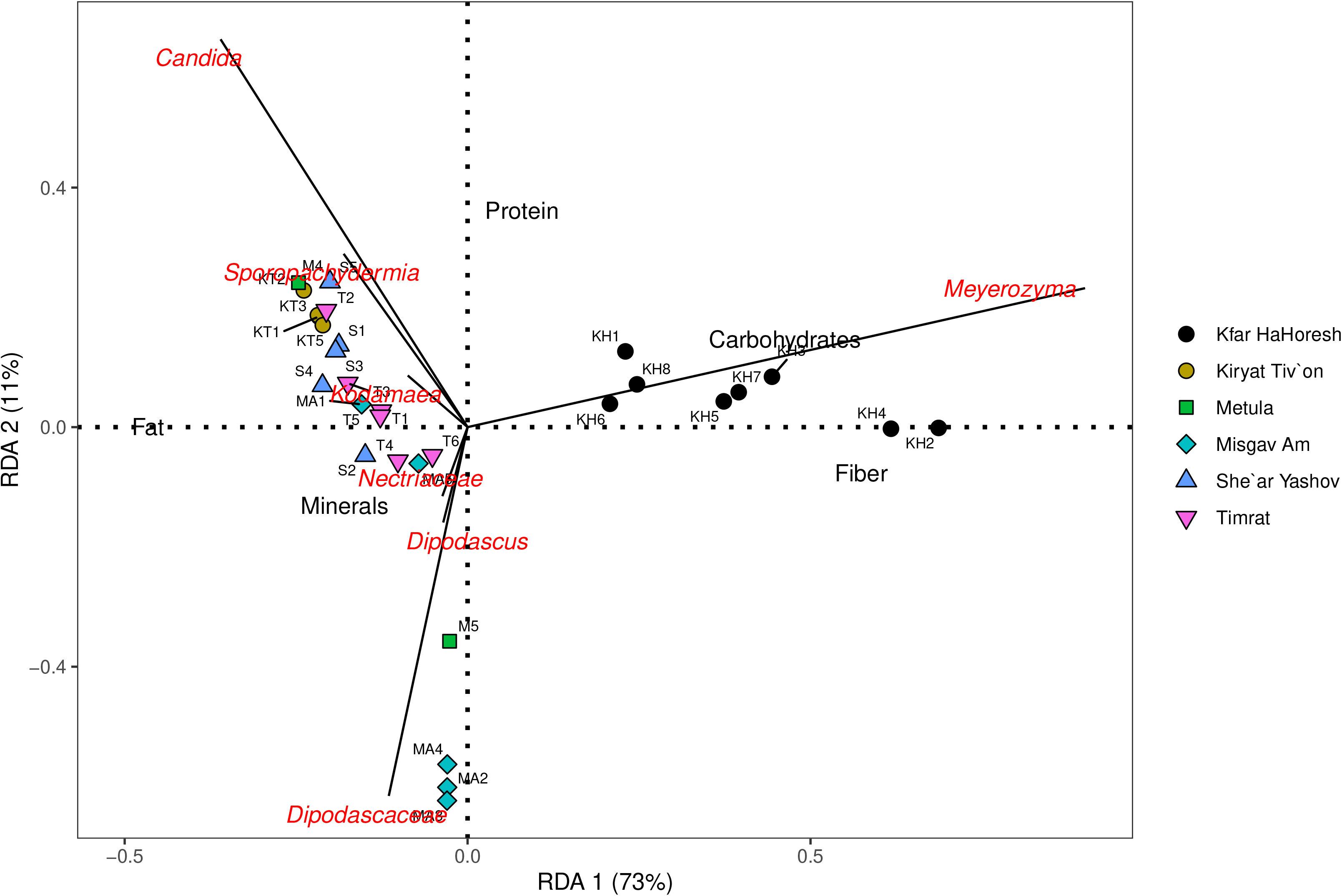
RDA of the fungal composition in the different locations and larval samples: KH - Kfar HaHoresh; KT - Kiryat Tiv’on; T – Timrat; MA - Misgav Am; M – Metula; S - She’ar Yashuv. The fungal genera and families that dominated the community composition are marked in red. The nutrient components of the compost are marked in black.

## DISCUSSION

The samples in this research were collected from various household composts in northern Israel. Although the microbial composition may change in the environment through time due to the composting process (17), the dominant fungal genus in most of the environments and larval guts was *Candida.* These results contradict previous analyses of the gut mycobiome composition of BSF larvae that were fed on agricultural waste (13) and of the mycobiome composition of the compost environment that was treated with BSF (18). In these cases, the most abundant fungal genus in BSF was *Pichia*, which was not an abundant genus in our study.

The mechanism behind the dominance of *Candida* in these environments is probably based on a fungal-insect interaction. There may be mutualistic interactions between the BSFL and *Candida*, as the larvae provide nutrients and mechanical defense for the fungi and the fungi provide supplementary metabolites or nutrients for the insects through direct consumption. Such mutualistc interactions are known from other dipterans, such as *Drosophila spp*., in which the fungus is known to affect insect survival and development time (19). *Candida* has also been found to be abundant in several *Drosophila* species (20). Conversely, the interaction may be commensalism, in which *Candida* does not provide any benefits for the insect, while the insect provides a vector for fungal dispersal to new resource patches and increases the probability of outbreeding (21). However, these dispersal “services” can have an indirect effect on the insect due to colonization of neutral fungi in the insect environment, which will compete with entomopathogenic fungi and reduce their ability to increase in population size (4) or due to secretion of mycotoxins that harm pathogenic fungi (13).

Given the fungal-insect interaction, we must also consider what determines the fungal composition of the insect environment and gut. *Candida spp*. may be common in the environment, and therefore, highly abundant in the insect gut. Alternatively, these species may be adapted to the insect gut and provide the insect with a benefit that allows the insect to disperse *Candida spp*. to new resource patches and increase their abundance in the environment. We must also determine whether the most important factor affecting fungal composition is the insect diet (22) or the environmental composition (23). It seems that in the two locations that were not dominated by *Candida*, KH and MA, the fungal community composition was influenced by the fiber content of the compost. The fungal composition in these locations was diverse without any dominant groups. However, the insect gut from these locations had higher abundance of *Meyerozyma* and Dipodascaceae (respectively), which were not abundant in the compost. The presence of these groups in the BSF gut was affected by the fiber and protein composition in the environment. Therefore, it seems that dominance of different species in the environment affects their dominance in the insect gut. However, in cases where dominant species are not present, the important factor determining the fungal composition of the insect gut is its diet. Moreover, consumption of a high fiber diet can increase the production of anti-microbial peptides (AMP) by the insect (24). These AMP are part of the insect’s immune system and can affect the microbial composition of the insect’s environment and gut. *Candida spp*., which were abundant at most sites, may be more susceptible to these AMP that are generated by diets high in fiber.

Understanding the metabolic effect of these fungi on the BSF larvae is still a black box. Little is known about the metabolic interactions of insects with eukaryotic microorganisms, such as yeast and yeast-like fungi and molds. As their metabolic complexity and ability is intense, they have the potential to dramatically affect the physiological condition of the insect. Revealing these metabolic interactions may help to increase the rearing efficiency of insects, such as the BSF, which is currently being reared at large scales as an alternative protein source (25).

## MATERIALS AND METHODS

### Larvae collecting

Stage 5 larvae were collected from household composts in Israel with similar waste characteristics (only vegetarian households) (Kiryat Tiv’on – KT (32°43’05’’ N 35°07’39’’ E); Timrat – T (32°42’12’’ N 35°13’31’’ E); Kfar HaHoresh – KH (32°42’06’’ N 35°16’24’’ E); Misgav Am – MA (33°14’51’’ N 35°32’54’’ E); Metula – M (33°16’38’’ N 35°34’41’’ E); She’ar Yashuv – S (33°13’36’’ N 35°38’47’’ E).). The larvae were placed in 95% ethanol upon collection and were kept in ice until storage at −20°C prior to further analysis.

### Nutrient analysis of the compost

In order to determine whether the yeasts in the environment were affected by the environmental conditions, the mega-nutrient composition of the compost was analyzed in three samples from each compost site. The protein content was measured using the Kjeldahl method for analyzing total nitrogen and calculating the protein ratio (14). Total lipid content was determined by extraction with hexane in a Soxhlet apparatus following the technique reported by Bligh and Dyer (15) with modifications. Fiber and mineral contents were measured after digestion with H_2_SO_4_ (1.25%) and NaOH (1.25%) and burning the remains at 600°C. Total carbohydrate content was calculated by subtracting the lipid, protein, mineral and fiber weights from the total dry weight (before the analyses were conducted).

### NGS of the mycobiome in the BSF gut and the substrate

Genomic DNA (gDNA) was extracted from BSF gut tissue using a Qiagen DNeasy blood and tissue kit. DNAseq libraries were produced according to the manufacturer’s protocol: “16S Metagenomic Sequencing Library Preparation protocol of Illumina”. The primers ITS1-ITS2 (ITS1: TCCGTAGGTGAACCTGCGG; ITS2: GCTGCGTTCTTCATCGATGC) were used in Illumina MiSeq, paired-end reads, 2×150bp. Demultiplexing and adaptor trimming was performed automatically on the MiSeq. Paired-end demultiplexed raw data reads were analyzed using the dada2 package in R (16). The taxonomic classification was conducted using a reference fungal dataset from the UNITE database site.

### Data analysis

Differences in the nutrient composition of the substrate were analyzed using Kruskal-Wallis non-parametric analysis. Similarities between the mycobiome composition of the larval gut and the substrate were analyzed on a proportion of the different genera using principal components analysis (PCA). The effect of substrate nutrient composition on community composition in the substrate and in the BSFL gut was tested using redundancy analysis (RDA). All statistical analyses were conducted using PAST and R software (16).

## ACKNOWLEDGMENTS

We would like to thank Llzar M. and Sherf G. for assisting with the NGS analysis. This research was support by the ISRAEL SCIENCE FOUNDATION (grant No.1167/21).

